# Defining the separation landscape of topological domains for decoding consensus domain organization of 3D genome

**DOI:** 10.1101/2022.08.08.503155

**Authors:** Dachang Dang, Shao-Wu Zhang, Ran Duan, Shihua Zhang

## Abstract

Topologically associating domains (TADs) have emerged as basic structural and functional units of genome organization, and have been determined by many computational methods from Hi-C contact maps. However, the TADs obtained by different methods vary greatly, which makes the accurate determination of TADs a challenging issue and hinders subsequent biological analyses about their organization and functions. Obvious inconsistencies among the TADs identified by different methods indeed make the statistical and biological properties of TADs overly depend on the method we chose rather than on the data. To this end, we employ the consensus structural information captured by these methods to define the TAD separation landscape for decoding consensus domain organization of the 3D genome. We demonstrate that the TAD separation landscape could be used to compare domain boundaries across multiple cell types for discovering conserved and divergent topological structures, decipher three types of boundary regions with diverse biological features, and identify <u>Cons</u>ensus <u>T</u>opological <u>A</u>ssociating <u>D</u>omains (ConsTADs). We illustrate that these analyses could deepen our understanding of the relationships between the topological domains and chromatin states, gene expression, and DNA replication timing. In short, we provide an alternative solution to deal with the serious inconsistencies of TADs obtained via different methods by defining the TAD separation landscape and ConsTAD.

## Introduction

Recent advances in imaging technologies (1, 2) and chromosome conformation capture (3C)-based technologies such as Hi-C (3–7) provide unprecedented opportunities to decipher the mysteries of three-dimensional (3D) genomic organizations in the nucleus of eukaryotes. It is gradually clear that the genome folds into a hierarchical configuration consisting of multi-scale chromatin structures such as chromosomal territories (6, 8, 9), A/B compartments (6, 7), topologically associating domains (TADs) (9, 10), and chromatin loops (7, 11). Among these structures, TADs, manifested as squares of increased intensity in the Hi-C contact maps, are structural domains with intensive self-interactions. They are demarcated by TAD boundaries (start and end regions of a domain), which serve as insulators to prevent inter-TAD interactions and favor intra-TAD interactions (9, 10). TADs are often considered as stable neighborhoods for gene regulation (7, 12, 13) and are reported to be highly conserved across multiple cell types and different species (7, 14–17). More importantly, many studies have shown that the disorder of TADs are closely related to some severe diseases (13, 18), even including cancer (19, 20). Therefore, TADs are the basic structural and functional units of chromatin organization, and the accurate determination of TADs is of vital importance for the 3D genome study.

In recent years, researchers have developed a variety of methods to define TADs based on diverse computational strategies (21–23). For example, some one-dimensional (1D) topological indicators including the directionality index (DI) (9), insulation score (IS) (10), and contrast index (CI) (24), were developed to capture the boundaries between adjacent TADs. TopDom (25), HiTAD (26), HiCDB (27) and OnTAD (28) were further designed based on some modified 1D indicators. Moreover, MSTD (29), IC-Finder (30), ClusterTAD (31) and CHDF (32) were designed to extract the interaction signals in the Hi-C contact map and then adopt some clustering algorithms to achieve it. Probabilistic models (e.g., GMAP (33) and HiCseg (34)) with certain assumptions have also been developed. Besides, some methods like 3DNetMod (35), Spectral (36) and deDoc (37) treated the Hi-C contact map as an adjacency matrix of chromatin interaction network and utilized community detection or graph segmentation algorithms for this task. However, different TAD-calling methods have identified very inconsistent and diverse TADs (21–23).

These TAD-calling methods were derived from a descriptive definition of TADs as squares with intensive self-interactions in the Hi-C contact maps. Due to the lack of a golden standard, it is difficult to interpret the inconsistencies among TADs from different approaches. Zufferey et al. systematically evaluated and compared some TAD-calling methods by designing a series of application scenarios and provided helpful guidelines for choosing an approach for a practical application (23). But the selected method cannot consider all aspects and may still suffer from certain limitations. Here, we apply 16 TAD-calling methods on Hi-C data generated for chromosome 2 of seven human cell lines (7). We find that different approaches returned quite diverse TADs when using the same Hi-C data, but each method could achieve more similar results even on Hi-C data of different cell lines, reflecting that these TADs were overly dependent on the method used rather than the data. Besides, through a boundary voting strategy, we reveal some distinctly inaccurate TAD boundaries in each method, while the consensus among different methods can guide us to find more reliable ones. Therefore, we define the TAD separation landscape based on the consensus structural information captured by different methods. The TAD separation landscape could display the boundary regions between TADs and serve as an indicator for boundary comparison to reveal the conserved and diverse boundaries across multiple cell types. Moreover, based on the boundary regions and ConsTADs revealed by the TAD separation landscape, we could decipher three types of boundary regions and five types of topological domains with various properties in terms of chromatin states, gene expression, and DNA replication timing. We expect this study can provide an alternative solution to determine accurate topological organizations of the 3D genome and characterize the biological properties of TADs.

## Results

### TADs identified by different methods show obvious inconsistency

We exemplify the diverse characteristics of TADs identified by 16 TAD-calling methods with the Hi-C contact matrices of GM12878 and K562 (**Fig. 1A and Fig. S1A**). TADs identified by different methods were quite diverse visually on the given data, but each method could get more similar TADs between GM12878 and K562 in this region. Besides, the measure of concordance (MoC) between TADs identified by different methods was smaller than that between TADs from the same method on two cell lines, which quantitatively suggests the inconsistency of TADs identified by different methods (**Fig. 1B**). We further confirmed that TADs found by different methods on the same data were more diverse in number and average size than those found by the same method across all the seven cell lines (**Figs. 1C, 1D and Figs. S1B, S1C**). Moreover, the TADs identified by the same method across cell lines could achieve larger consistency in terms of two metrics compared with those obtained by different methods on the same data (**Figs. 1E, 1F**). We can observe similar results for all the TAD-calling methods (**Figs. S1D-G**). These results suggest that the existing methods could identify very inconsistent TADs and the TADs heavily depend on the method used rather than the difference of Hi-C data, which might obscure the characterization of real topological domains.

**Figure 1.**
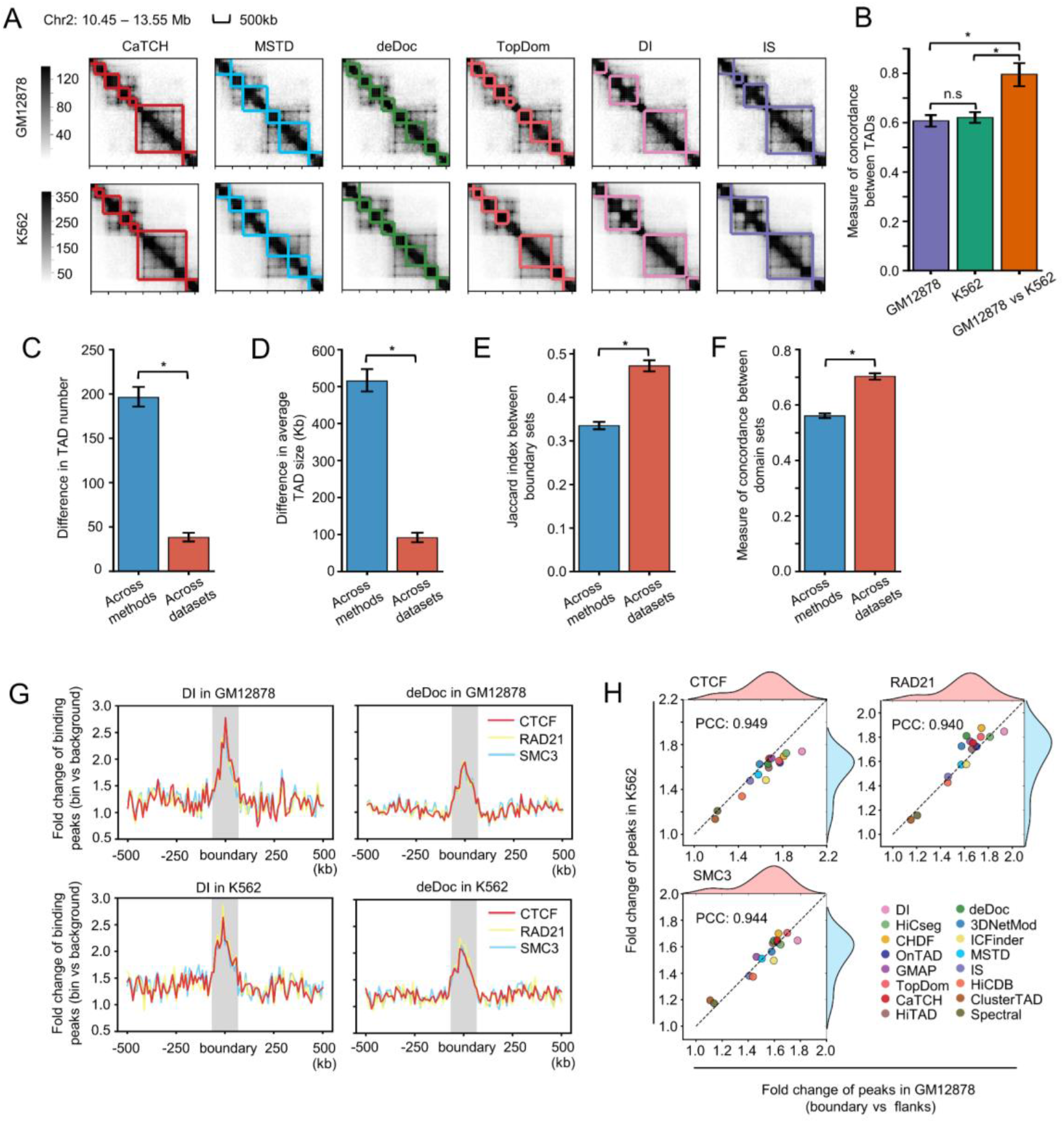
Comparison of TADs identified by 16 TAD-calling methods on Hi-C data from diverse cell lines. (A) TADs identified by different methods on the same chromatin region of GM12878 and K562. The results of six methods are shown and the remaining ones are shown in Fig. S1A. (B) Comparison of measure of concordance (MoC) between TADs identified by different methods in GM12878 or K562, and between TADs identified by the same method in two cell lines, for the regions in (**A**). (C-F) Comparison of the difference in TAD number (C), TAD size (D), the Jaccard index between boundary sets (E) and the MoC between domain sets (F) across TADs identified by different methods on the same dataset or across TADs identified by the same method on different datasets. The error bars represent the 95% confidence intervals. Mann-Whitney U tests are performed and * represents p-value < 0.0001. (G) Average profiles for fold changes of CTCF, RAD21 and SMC3 binding peaks around boundaries identified by DI or deDoc in GM12878 and K562. The fold changes are calculated for each 10kb bin by dividing the expected peak numbers in chromosome. (H) Comparison of fold changes of CTCF, RAD21 and SMC3 binding peaks for boundaries identified by the 16 methods between GM12878 and K562. The fold change for each method is calculated by dividing the signal of the boundary by the average signal of flanking regions in the profiles shown in figure G. The distribution of fold changes for different methods in GM12878 or K562 are shown and their Pearson correlation coefficient (PCC) are calculated.

More importantly, the inconsistencies among TADs identified by different methods raise our concerns about the subsequent biological analyses that rely on these TADs. For instance, the cohesin complex and chromatin insulator protein CTCF are frequently reported to enrich in TAD boundaries, and they appear to be necessary for boundary formation, maintenance and remodeling (7, 38–40). Thus, we tested whether the TAD boundaries identified by different methods (e.g., DI and deDoc) were enriched with CTCF and two main components of cohesion complex, RAD21 and SMC3, in GM12878 and K562. All three proteins were enriched in the TAD boundaries from TADs of both methods, but the significance levels of enrichment were different. The DI boundaries showed stronger enrichment in both GM12878 and K562 than those by deDoc (**Fig. 1G**). We further observed that the enrichment of three proteins in TAD boundaries varied a lot among multiple methods, but the same method could achieve similarly results between two different cell lines (**Fig. 1H**). Thus, performing biological analysis based on TADs derived from different methods could result in quite different observations.

### Boundary voting reveals the unreliability of TADs of individual methods

Here, we evaluate TADs from different methods based on a boundary voting strategy (**Fig. S2A**), and assign a boundary score to each bin along the chromosome by counting the number of methods that define it as a TAD boundary. Thus, the boundary scores are integers ranging from 0 to 16. The distribution of boundary scores for bins on chromosome 2 of GM12878 DpnII replicate indicates the presence of many low-scoring boundaries (**Fig. S2B**). By dividing the boundary scores into five intervals, we found that each method could return some low-scoring boundaries with a score of 1 or 2, but the proportions varied across methods (**Fig. 2A**). For example, OnTAD and DI possessed significantly fewer low-scoring boundaries, while Spectral and HiCDB had a much higher proportion of them. More importantly, boundaries belonging to different score intervals show diverse profiles for indicators including DI, IS and CI and different enrichment patterns for three structural proteins. We can observe that the higher-scoring boundaries show stronger insulation strength of chromatin interactions as well as higher level of enrichment for CTCF and cohesin complex (**Fig. 2B**). On the contrary, boundaries belonging to the score interval of 1 to 2 showed completely counterintuitive patterns for both 1D indicators and structural proteins, indicating these boundaries were not reliable. Besides, after collecting all candidate boundaries (bins with non-zero scores) found by the 16 methods, we find that some methods could return more potential boundaries, but would also bring in more low-scoring ones, such as HiCDB and deDoc, while some methods like DI and OnTAD tend to capture boundaries with higher scores to ensure their reliability, but may miss more candidate ones with moderate scores (**Fig. 2C**). By computing the minimum distance between non-zero scoring bins and the boundaries returned by each method, we observe that there are always some high-scoring bins located far away from the boundaries, which is very prominent for methods like DI and OnTAD (**Fig. S2C**).

**Figure 2.**
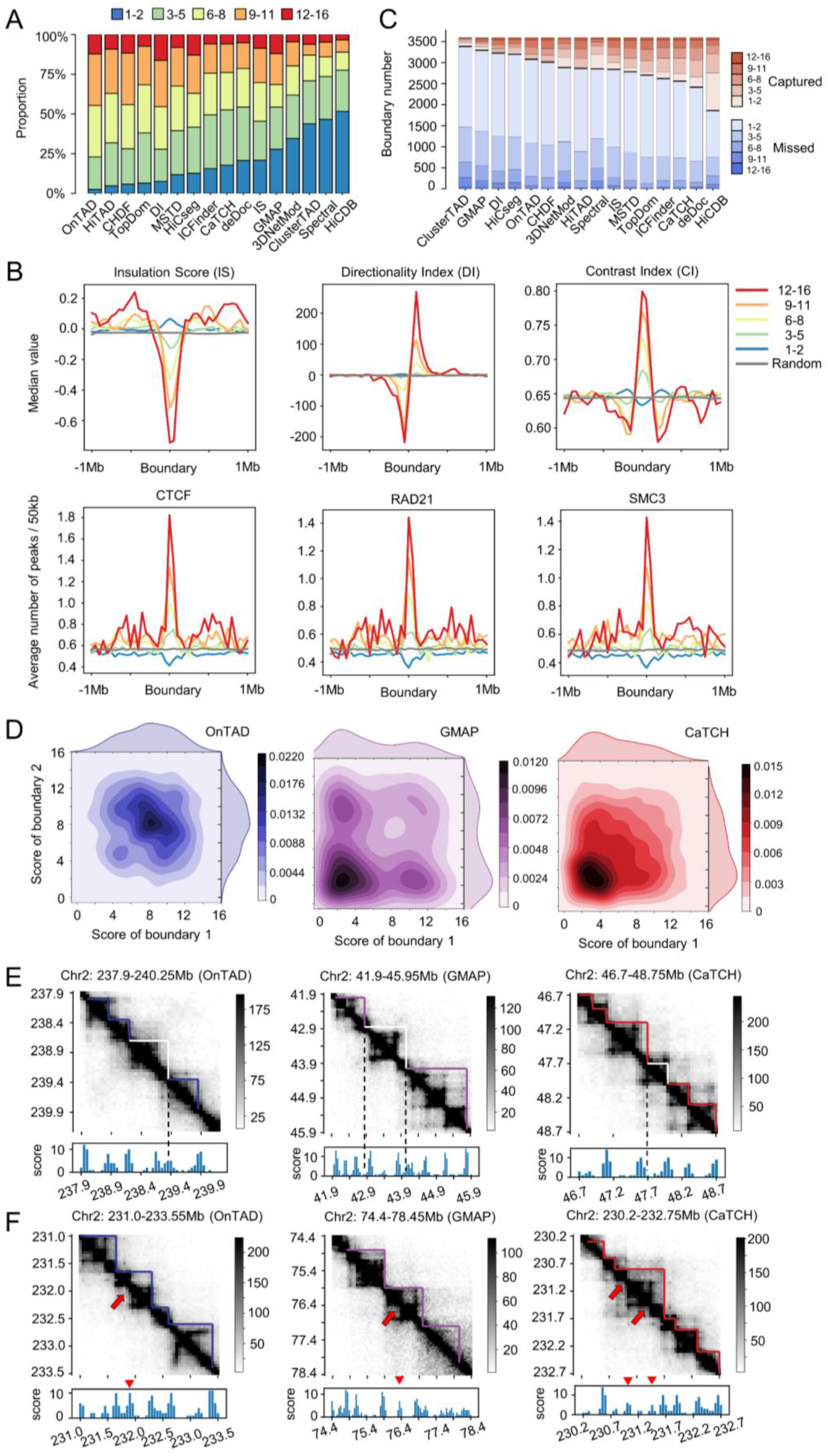
Systematic evaluation of 16 TAD-calling methods based on the boundary voting strategy. (A) Proportion of boundaries with different levels of boundary score for all 16 methods. These methods are sorted in ascending order by the proportion of boundaries belonging to the first level (ranging from 1 to 2). (B) Profiles of three topological indicators (Insulation Score [IS], Directionality Index [DI], Contrast Index [CI]) and profiles of three structural proteins (CTCF, RAD21, SMC3) within 2Mb regions centered on boundaries with different boundary score levels or randomly selected regions. (C) Number of boundaries with different boundary scores captured or missed by each method. These methods are sorted in ascending order by the number of captured boundaries. (D) Density profiles of two boundary scores for TADs identified by different methods. The results of three representative methods including OnTAD, GMAP and CaTCH are shown and results of other methods are shown in Fig. S3. (E) Hi-C contact maps around the unreliable TADs with low boundary scores for three representative methods including OnTAD, GMAP and CaTCH. The unreliable TADs are indicated by white frames and the low-scoring boundaries are indicated by dashed lines. (F) Hi-C contact maps around the high scoring boundaries that missed by certain method. The examples for three representative methods including OnTAD, GMAP and CaTCH are shown. The missed boundaries are indicated by red arrows and the corresponding boundary scores are indicated by red triangles.

More interestingly, the paired score distribution of two boundaries of individual TADs returned by each method demonstrate three main types of patterns (**Fig. 2D** and **Fig. S3**). For OnTAD, the paired scores are relatively high and similar, making the density center of the distribution close to upper right. For GMAP, the paired scores show a clear imbalance, where one boundary of a TAD has a high score and the other shows a lower one, reflecting that one boundary of the TAD tends to be not reliable enough. For CaTCH, the scores of both boundaries for most TADs were low, making the density center of the distribution close to the lower left, raising our concerns about the reliability of these TADs. Based on these paired scores for TADs, we could discover some unreliable domains and missed boundaries for each method (**Figs. 2E, 2F** and **Fig. S4**). It is clear that the unreliable TADs with low-scoring boundaries are visually inconsistent with the domain patterns exhibited in the Hi-C contact maps (**Fig. 2E**) and the missed boundaries could match well with some peaks in the boundary score profile (**Fig. 2F**). All these results indicate that existing TAD-calling methods have distinct limitation, but the consensus among them could lead us to find more reliable TADs.

### Define the TAD separation landscape based on consensus of all TAD-calling methods

To overcome the drawbacks of individual TAD-calling methods and well employ the consensus structural information captured by these methods, we suggest a computational strategy to dissect and integrate the TADs identified by existing methods. First, we apply all TAD-calling methods to the same Hi-C contact map and collect the identified TAD boundaries. Second, we construct the boundary score profile of all bins along the genome based on the boundary voting strategy. Third, we refine the boundary score profile through three kinds of additional operations to get the TAD separation landscape (**Fig. 3A**).

**Figure 3.**
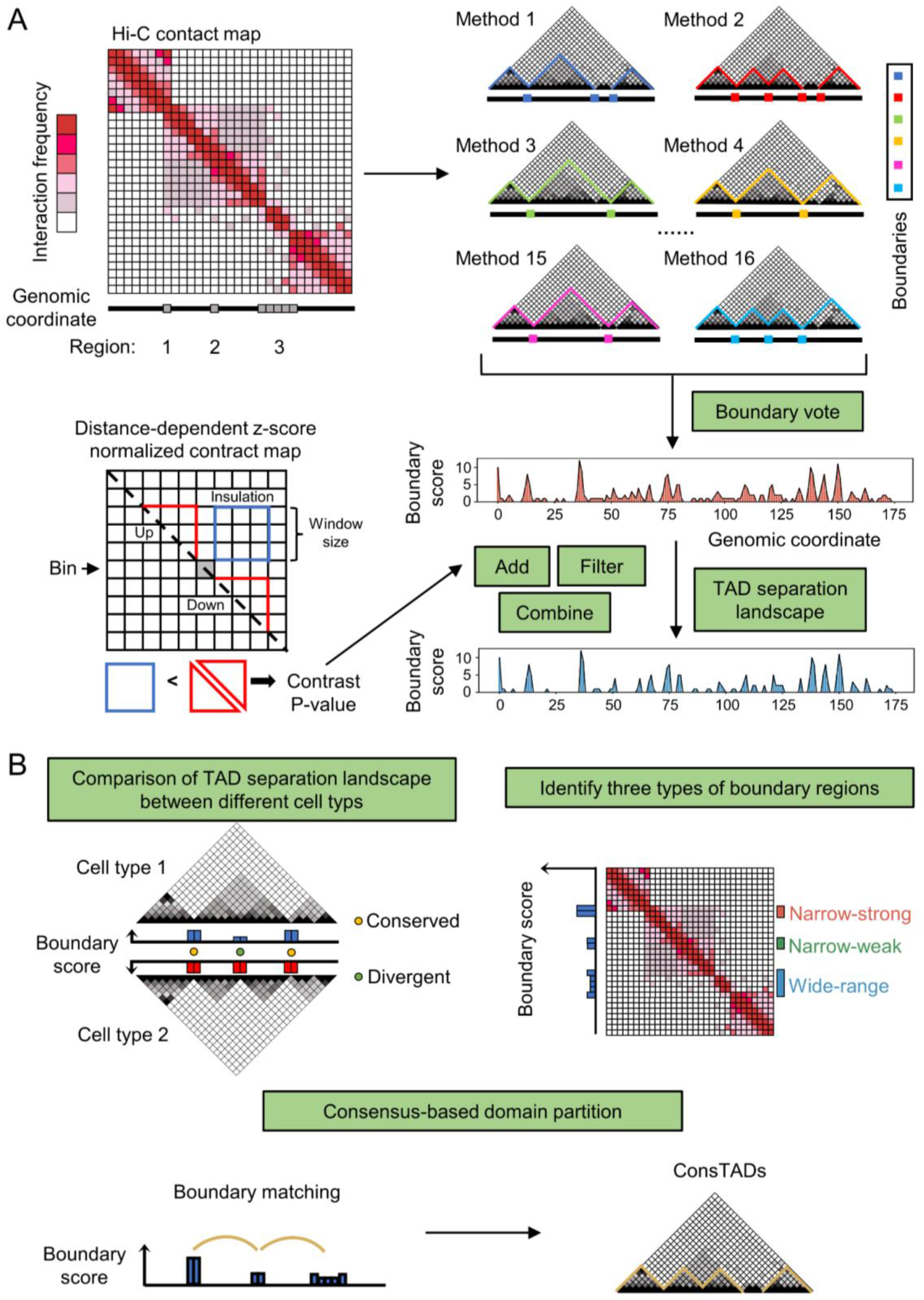
Overview of the construction of the TAD separation landscape and its three applications. (A) The representative Hi-C contact map contains several candidate boundary regions and different TAD-calling methods identify diverse TADs with variable boundaries. Based on the boundary voting strategy, each bin on the genome will get a score to indicate how many methods consider it to be a TAD boundary. The TAD separation landscape is constructed from the original boundary score profile using three additional operations, including **Add**, **Filter** and **Combine**, which are calculated based on the contrast *p*-value, an indicator to reflect the insulation effect of chromatin interactions for each bin. (B) Three applications of the TAD separation landscape including domain boundary comparison, boundary type identification and consensus domain detection.

Intuitively, the consecutive bins with non-zero scores represent the potential boundary regions. However, the original boundary score profile contains numerous low-scoring bins, which are unreliable and noisy and make the adjacent boundary regions too close to each other (**Figs. S5D-F**). Therefore, we introduced an additional 1D indicator, called the contrast *p*-value, to refine the boundary score profile. For each bin, the contrast p-value reflects the degree of difference between the chromatin interactions within the upstream and downstream regions and the interactions between them (**Fig. 3A**). Next, three kinds of operations are performed to modify the boundary score profile and construct the TAD separation landscape based on these contrast p-values (see **Methods**). We tested these operations on chromosome 2 of GM12878 MboI replicate and found that they could filter out some unreliable bins with low scores (**Fig. S5E**) and adjust the distance between adjacent TAD boundaries (**Fig. S5F**). Experiments in other cell lines further demonstrate that the refined profiles possess boundary regions with higher scores, and the distances between adjacent regions become larger and more consistent with the lengths of topological domains identified by general TAD-calling methods (**Figs. S5G-J**). We refer to the refined boundary score profile as the TAD separation landscape, which integrates the results of a set of TAD-calling methods and can precisely depict the locations of potential boundary regions between adjacent TADs (**Fig. 3A**).

The TAD separation landscape could be used as an effective indicator for comparing topological domains cross cell types to reveal the conserved and cell type-specific boundaries between different cell lines. It could indicate the presence of three different types of boundary regions with distinct chromatin insulation patterns and biological functions. More importantly, it could contribute to the detection of consensus topological associating domains (ConsTADs) for identifying more accurate domains (**Fig. 3B**).

### Comparing TAD boundaries cross cell types based on the TAD separation landscape

We calculated the corresponding TAD separation landscapes for the MboI Hi-C contact maps of chromosome 2 in seven human cell lines, respectively. The boundary regions indicated by the TAD separation landscapes matched well with the domain borders in the Hi-C contact maps (**Fig. 4A**). Then we divide all bins along the chromosome into six levels according to their average boundary scores across seven cell lines. The higher the level, the larger the boundary score and the first level only contains bins with a score of 0. We find that bins with higher boundary score level are more conserved across cell types and enriched for more housekeeping genes as well as CTCF binding peaks, and the regions bounded by CTCF show higher level of cross-species sequence conservation (**Fig. 4B**). In addition, the clustering results of multiple Hi-C samples based on their TAD separation landscapes are well consistent with organogenesis, with cell lines originating from the same germ layer being clustered together (**Fig. 4C**). Therefore, these results suggest that the TAD separation landscape can precisely characterize the locations of TAD boundaries and reflect their biological significance and cell-type specificity..

**Figure 4.**
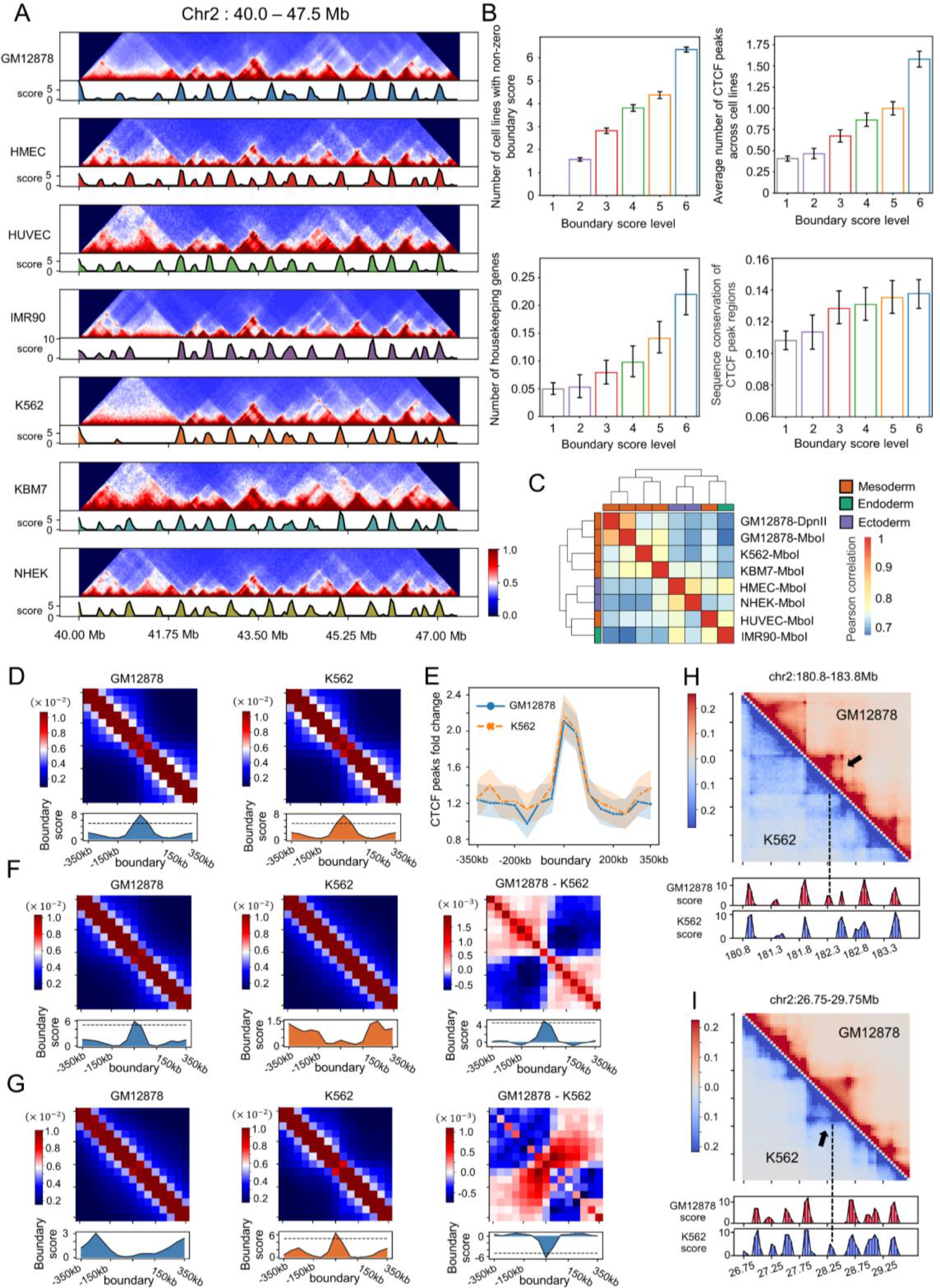
Comparison of TAD boundaries across different cell lines. (A) Hi-C contact maps and the corresponding TAD separation landscapes of a chromatin region (chr2:40.0-47.5Mb) in seven cell lines. (B) The relationship between the boundary score level and the conservation of boundary across cell lines (upper left), the average number of CTCF peaks across cell lines (upper right), the number of housekeeping genes (bottom left) and the conservation of DNA sequence for CTCF binding regions (bottom right) respectively. (C) Clustering of multiple Hi-C samples from different cell lines based on their TAD separation landscapes. (D) Aggregated Hi-C contact maps around the conserved boundary regions between GM12878 and K562, combined with the average boundary score profiles of the corresponding regions. The dotted lines indicate the boundary score of five. (E) Profiles of CTCF peak fold change (bin vs background) around the conserved boundary regions between GM12878 and K562. The shaded areas represent the 95% confidence intervals in 1000 boostraps. (F and G) The aggregated Hi-C contact maps around the GM12878-gained boundary regions (F) and GM12878-lost boundary regions (G) in GM12878 and K562 cell lines, combined with the average boundary score profiles of the corresponding regions (left and middle panel). The difference between these aggregated maps and the average boundary scores (right panel). (H and I) The contrastive Hi-C contact maps around a GM12878-gained boundary region (H) and a GM12878-lost boundary region (I), compared with K562. The Hi-C contact maps and corresponding boundary score profiles in GM12878 (marked in red) and K562 (marked bin blue) are shown. The dashed lines and arrows indicate the location of the boundary regions.

Moreover, the TAD separation landscapes could not only indicate the locations of TAD boundaries, but also reflect the conservatism and heterogeneity of topological organizations among different cell lines (**Fig. 4A**). Taking the comparison of GM12878 and K562 as an example, we showed that the TAD separation landscapes could serve as effective indicators to facilitate the identification of conserved and cell type-specific TAD boundaries between them. As shown in **Fig. 4D**, the aggregated Hi-C contact maps around the conserved boundary regions between GM12878 and K562 exhibit significant insulation of chromatin interactions, coupled with the peaks of average boundary scores in both cell lines. These conserved boundaries show very similar patterns of CTCF enrichment in both cell lines (**Fig. 4E**). In addition, according to the difference in the TAD separation landscapes, we could identify some GM12878-gained boundaries, which showed insulation of chromatin interactions and peaks of boundary score only in GM12878, while no evident insulation and boundary pattern was seen in K562. When we subtracted the Hi-C contact map of K562 from the contact map of GM12878, the difference between them became more obvious. The upstream and downstream regions of the boundary show positive signals, while the boundary regions show negative ones, indicating stronger insulation of chromatin interactions in GM12878. Consistent with this, subtracting the boundary score profile of K562 from the profile of GM12878 still presented a peak at the boundary (**Fig. 4F**). In contrast, the GM12878-lost boundaries show completely opposite patterns. They don’t behave like TAD boundaries in GM12878, but show stronger insulation of chromatin interactions and larger boundary scores in K562 (**Fig. 4G**). However, close to the GM12878-gained and GM12878-lost boundaries, we cannot see significant difference between the CTCF binding profiles of GM12878 and K562 (**Fig. S6B**). This is probably because the cell type-specific boundaries are less dependent on CTCF, which has also been reported in a recent study (16). For the GM12878-gained and lost boundary regions when compared with K562, we also observe that the TAD separation landscapes display distinct changes between the topological domains in these cell lines (**Figs. 4H, 4I**). In addition to the comparison between GM12878 and K562, we also performed pairwise comparisons between seven cell lines based on their TAD separation landscapes for the conserved, cell type-gained and cell type-lost boundaries (**Figs. S6C, S6D**). The cell type-specific boundaries only occupied a relatively small part, and nearly half of the boundaries were conserved between two cell lines, confirming that TADs are highly conserved among cell types (7, 9).

Apart from the pairwise comparisons between cell lines, the TAD separation landscapes could also facilitate the boundary comparison among multiple cell types. Using the boundary score as an indicator, we can identify the conserved boundaries shared by all seven cell lines and the specific boundaries unique to each cell line (**Fig. S6E**). The conserved boundaries exhibit strong insulation of chromatin interactions and boundary score peaks in all cell lines, while the cell type-specific boundaries only show these patterns in the corresponding cell line (**Figs. S7A, S7B**). In addition to the most conserved and specific boundaries, the TAD separation landscapes can also reveal boundaries shared by part of the cell lines (**Fig. S8**), which match well with the domain borders in the corresponding Hi-C contact maps. These results suggest that the TAD separation landscapes could capture both the commonalities and differences in the topological organizations between different cell types.

### TAD separation landscape divides boundary regions into three different types with strong biological relevance

The boundaries revealed in the TAD separation landscape are variable in length and boundary scores. Based on the number of bins contained in each region and the average boundary score of them, we divided the boundary regions into three different types: narrow-strong boundary regions (NSBs), narrow-weak boundary regions (NWBs) and wide boundary regions (WBs) (**Fig. 5A**). The NSBs and NWBs typically cover few bins but with high and low boundary scores respectively, while the WBs can occupy many bins with moderate scores. The aggregated Hi-C contact maps around three different types of boundary regions indicate that they have distinct insulation patterns of chromatin interactions. The NSBs show strong insulation against chromatin interactions, whereas the NWBs exhibit local insulation with weaker strength and the WBs are larger fragments separating two adjacent domains. The insulation patterns of these boundary regions can be well captured by the boundary score profiles (**Fig. 5B**). We identified all three types of boundary regions in seven cell lines (**Fig. S9**) and in each cell line, the NWBs always account for the largest proportion, while the NSBs and WBs were slightly fewer (**Fig. 5C**). However, the cross-cell types comparison reveals that the NWBs are less conserved among different cell lines, while the NSBs exhibit a high degree of conservation, implying that different cell lines possess similar strong insulation patterns near these regions (**Fig. 5D**). Coherently, the WBs exhibit wide-range insulation patterns as well as broad distributions of protein binding peaks and the NSBs show sharp insulation patterns and concentrated protein binding profiles, while the NWBs have moderate and localized insulation effect and are less dependent on the structural proteins (**Figs. 5E, 5F**).

**Figure 5.**
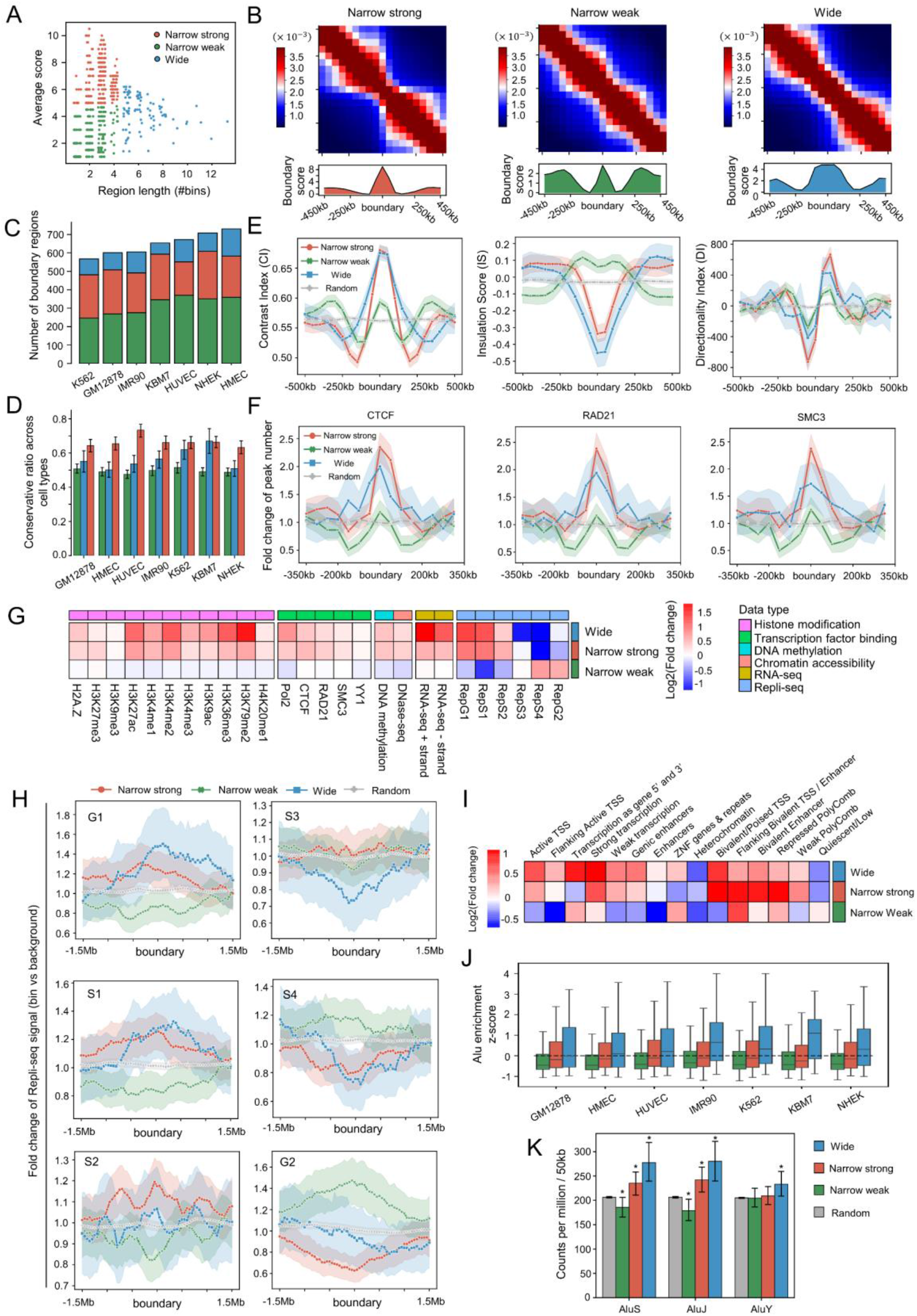
Three different types of boundary regions defined based on the TAD separation landscape. (A) Scatter plot of boundary regions based on their lengths and average boundary scores. (B) Aggregated Hi-C contact maps around NSBs (left panel), NWBs (middle panel), WBs (right panel), combined with the average boundary score profiles in GM12878. (C) Number of three types of boundary regions identified in seven cell lines respectively. (D) Conservation across cell lines of three types of boundary regions in seven cell lines. (E) Profiles of three kinds of numerical indices including CI (left panel), IS (middle panel), DI (right panel) around different types of boundaries and random control in GM12878. The shaded areas represent the 95% confidence intervals in 1000 boostraps. (F) Profiles of three kinds of biological signals including CTCF (left panel), RAD21 (middle panel), SMC3 (right panel) around different types of boundaries and random control in GM12878. (G) Fold change profiles of multiple types of biological data constructed for three types of boundary regions in GM12878. Fold change of a biological mark in each type is defined as the median signal of the mark in boundary regions divided by the median signal across the whole chromosome. (H) Fold change profiles of Repli-seq signal around three types of boundaries in six different phases of cell cycle: G1, S1, S2, S3, S4, and G2. (I) Fold change profiles of ChromHMM states constructed for three types of boundary regions in GM12878. The fold change of a chromatin state in each type is defined as the total length of the state in boundary regions divided by the state length across the whole chromosome. (J) Enrichment of Alu elements in three types of boundaries for seven cell lines. (K) Density of the Alu subclasses in three types of boundaries and random selected regions in GM12878. Mann-Whitney U tests were performed between three types of boundary regions and random regions, respectively, * represents *p*-value < 0.01.

Moreover, the WBs are enriched with active histone modifications such as H3K27ac, H3K36me3 and H3K79me2, and they also have stronger RNA polymerase II (Pol2) signals and chromatin openness, exhibiting significantly strong transcription (**Fig. 5G**). While the NSBs showed similar enrichment patterns but with weaker intensity (**Fig. 5G**, **Fig. S12A**). The NWBs do not show significant enrichment in most biological data, but possess distinct patterns on the Repli-seq signals that are entirely different from the WBs and NSBs (**Fig. 5G**). Repli-seq maps the sequences of nascent DNA replication strands throughout the whole genome during each of the six cell cycle phases, quantifying how early or late DNA replication occurs (44). Based on the enrichment profiles of Repli-seq signals around the three types of boundary regions during the six phases of the cell cycle, we observe a gradual shift of the replication patterns, in which the WBs and NSBs show signal enrichments in some early stages, like G1, S1 and S2, but decrease in some later stages, such as S3, S4 and G2. In contrast, the NWBs show opposite trends and the nascent DNA replication strands remain low level in the early phases but tend to be enriched in the later phases (**Fig. 5H**). These results suggest that the DNA sequences near the WBs and NSBs tend to replicate earlier than sequences near the NWBs (**Fig. S13B**). Based on the enrichment analysis of 15 chromatin states annotated by ChromHMM (45) in three types of boundary regions of GM12878, we observe that the WBs are enriched with Active TSS, Strong transcription and some other chromatin states associated with active gene transcription, and NSBs are enriched with stronger enrichment of bivalent chromatin regions, such as Bivalent TSS and Bivalent enhancer as well as polycomb group of proteins (**Fig. 5I**). We also observed similar patterns in other cell lines (**Fig. S10A**). Bivalent chromatin segments have been found to play vital roles in the developmental regulation of pluripotent embryonic stems cells, as well as in gene imprinting and some diseases (46–50). Recalling the strong conservation of NSBs across cell types (**Fig. 5D**) and the general enrichment of bivalent chromatin segments on these boundary regions in multiple cell lines (**Fig. S10A**), we guess that the NSBs might be crucial for stem cell differentiation and cell fate decision, which deserve further exploration. We also explored the relationships between these boundary regions and the chromatin states annotated by Segway-GBR (51) as well as another chromatin structures termed as subcompartments (7, 52) (**Figs. S11B, S11C**). The WBs are highly related to both broad and cell type-specific gene expression and the NSBs are associated with broad gene expression and the facultative heterochromatin, while the NWBs typically belong to the constitutive heterochromatin or quiescent chromatin regions. These results are in accordance with the assignment of subcompartments to these boundary regions. The WBs mainly belong to the A1 subcompartments that have been reported as transcriptionally active regions and the NSBs are enriched with A2 subcompartments, which were considered as active regions resembling the A1 type, but in a recent study, they were indicated to be intermediate subcompartments containing some poised promoters (53). While the NWBs are mainly divided into B-type subcompartments, which usually serve as transcriptionally repressed regions.

Moreover, for two types of repeat elements, Alu and TcMar-Tigger, relating to 3D chromatin architectures and regulatory elements (54, 55). a gradual increasing enrichment can be seen from two types of narrow boundary regions to the WBs, which exhibit stronger enrichment of both repeat elements relative to the random control in multiple cell lines (**Fig. 5J** and **Figs. S11A-S11G**). Both WBs and NSBs are enriched with AluS and AluJ compared to the random ones, while for the younger AluY, only the WBs exhibit a significant enrichment (**Fig. 5K**). Our results are consistent with some previous studies, reporting the enrichment of Alu elements in TAD boundaries (9, 15), and further suggest that different types of boundary regions may be enriched with certain sub-class of Alu. Moreover, we also observe a clear enrichment of another transposable element called TcMar-Tigger, which was rarely reported to be associated with TAD boundary and its function need to be further investigated. Furthermore, we checked the DNase-seq peaks in these boundary regions from GM12878, for enrichment of accessible motif instances of transcription factors (TFs) from HOMER database (56) (**Fig. S12C**). For all three types of boundary regions, some common TFs like CTCF and CTCFL are shared, and some special TFs for each type can also be found, such as Tcf12 for NSBs, SpiB for NWBs and NeuroD1 for WBs in GM12878, which might promote the study and understanding of the formation and maintenance mechanisms of these boundary regions in certain cell types (**Figs. S12**, **S13**).

### ConsTADs facilitate the understanding of biological modifications and DNA replication patterns within topological domains

Based on the TAD separation landscape, we design a boundary matching strategy to identify the ConsTADs to represent the topological domains under the consensus of the 16 TAD-calling methods. We first explored the profiles of some TFs and epigenomic modifications within these ConsTADs in GM12878 and found two kinds of distribution patterns. First, some TFs like CTCF, SMC3, RAD21 as well as the chromatin accessibility are more enriched or stronger in boundary regions than domain interiors, which is consistent with the conventional observation that CTCF and cohesins could contribute to the formation and maintenance of topological domain through binding at the boundary regions (7, 38–40). Second, some histone modifications such as H3K27ac, H3K4me3 and H3K79me2 are more abundant inside the domains, probably because more regulatory events are involved in domain interiors and therefore the modifications associated with the activity of regulatory elements such as enhancers and promoters are stronger within domains (**Fig. 6A and Fig. S15B**). In addition, the ConsTADs could capture more topological domains with significant H3K36me3 or H3K27me3 differential signals compared to the 16 TAD-calling methods (**Fig. 6B**) and the switching points between the H3K36me3-biased regions and H3K27me3-biased regions are more closer to the boundaries of ConsTADs than those of most other methods, except for HiCDB (**Fig. S15C**), which gave more boundaries and many of them had a low boundary score (**Fig. 2B**).

**Figure 6.**
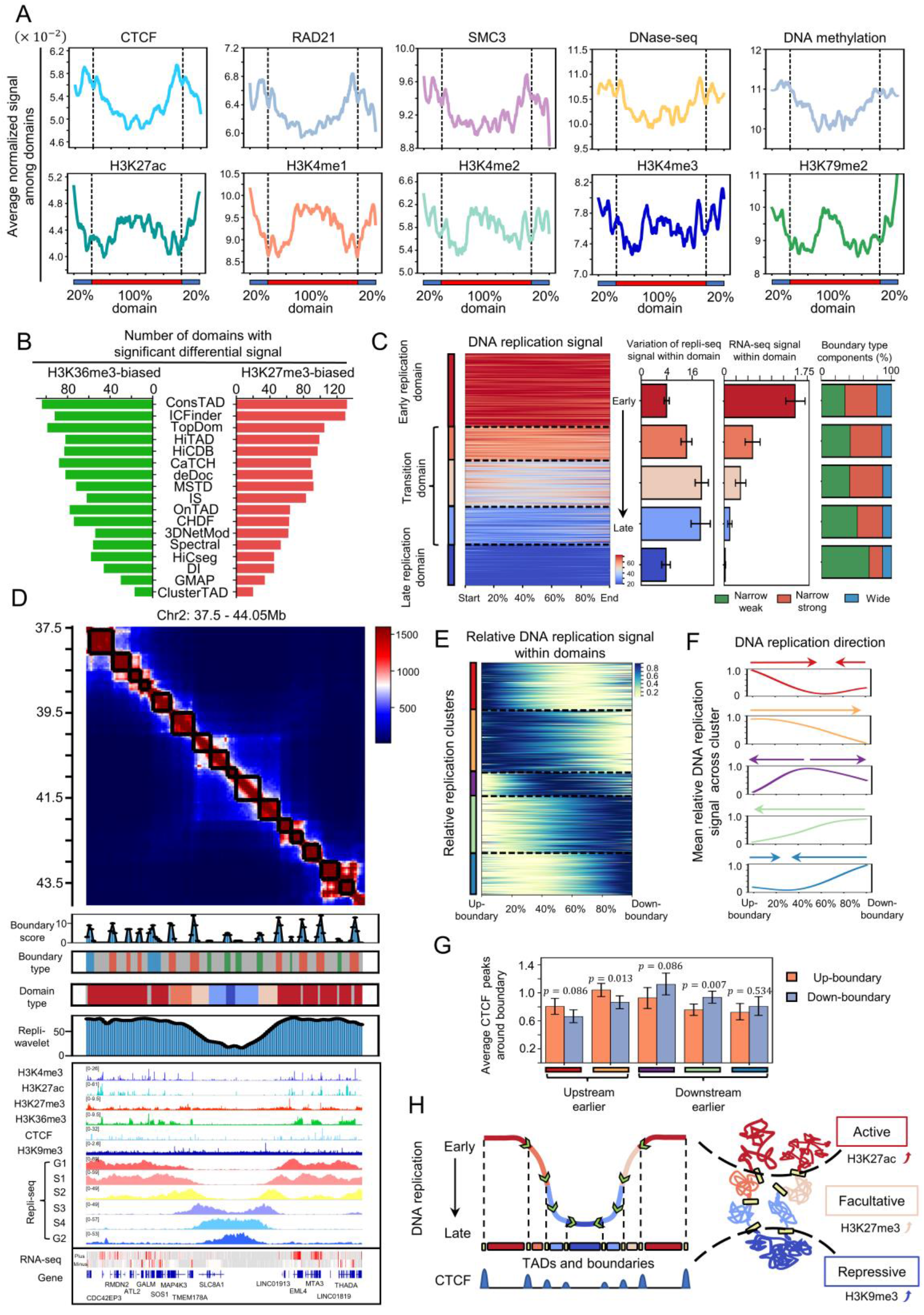
Biological relevance of ConsTADs. (A) Relative profiles of several biological features within ConsTADs and adjacent regions in GM12878. (B) Number of domains with significant differential signal of H3K36me3 or H3K27me3 from ConsTADs and the 16 TAD-calling methods. (C) Five types of ConsTADs with distinct DNA replication signals accompanied by the variances of Repli-seq signal, the average RNA-seq signal and the boundary type components within each type of domains. (D) Hi-C contact maps of a region on chromosome 2 of GM12878 accompanied by the ConsTADs (black frames in contact map) and the corresponding domain types identified in (**C**). (E) Five clusters of ConsTADs with distinct relative DNA replication models, which indicate the relative early or late replication times of chromatin regions within each domain. (F) Mean profiles of relative DNA replication signals for five clusters of ConsTADs in (**E**). Arrows indicate the direction of DNA replication from early to late. (G) Average number of CTCF binding peaks around the upstream and downstream boundary for each domain in five clusters. Mann-Whitney U tests were performed to get the p-values. (H) The replication domain model. Left, DNA replication timing across TADs belonging to early, transition and late replication domains accompanied with the CTCF profiles at TAD boundaries. Right, the corresponding chromatin arrangement of replication domains and their chromatin states and prominent modifications.

In terms of DNA replication, Pope et al. found that the human genome could be divided into some constant timing regions (CTRs) with relatively uniform replication timing, while the early and late CTRs were interrupted by some timing transition regions (TTRs). Based on the TADs identified by DI, they concluded that TAD boundaries usually isolated the early CTRs from TTRs, whereas the TTRs and neighboring late CTRs predominantly belonged to the same TADs (57). However, based on our analysis, DI tends to identify a small number of TADs with relatively large scale and may lack a fine depiction of the topological domains (**Figs. S1A, S1B**). Here, based on the Repli-seq data of GM12878, we divided ConsTADs into five clusters containing clusters of early and late replication domains, as well as three clusters of transition domains with replication timing in between (**Fig. 6C and Fig. S15D**). We could observe relatively uniform replication signals within the early and late replication domains, while the transition domains show larger variances, indicating a gradual DNA replication timing within them. The early and late replication domains perhaps correspond to the early and late CTRs that Pope et al. found, and benefiting from the intensive depiction of topological domains by ConsTADs, we found the timing transition regions might also correspond to some independent TADs, rather than being contained in the same TADs with late CTRs (**Fig. 6C**). Compared the TADs identified by the 16 TAD-calling methods, the replication domains obtained by ConsTADs had the highest approval degree among the results of all methods (**Figs. S15E-S15G**).

Besides, the five clusters of replication domains obtained from ConsTADs show different epigenomic patterns and gene expression levels (**Fig. 6C and Fig. S16A**). Some active markers, such as H3K4me3, H3K27ac and H3K36me3, as well as the gene expression, are the strongest in early replication domains and get weaker in three types of transition domains sequentially as the timing of replication changed from early to late, and become the weakest in the late replication domains. The repressive marker H3K9me3 show an opposite pattern with progressively stronger signal from early replication domains to the late ones, while the facultative marker H3K27me3 was stronger in three types of transition domains (**Fig. S16A**). It is known that both H3K9me3 and H3K27me3 are markers associated with gene repression, but unlike H3K9me3, which keeps genes silenced all the time and prevents binding of multiple TFs, H3K27me3 still allows genes to be activated through TF binding in different cell states (50, 58). These results suggest that the topological domains not only have different DNA replication timings, but might also correspond to different chromatin states. Correspondingly, we found that the early replication domains are typically enriched in active subcompartments like A1 and A2, and belong to regions with strong transcriptional activity, while the late replication ones mainly fall in the quiescent regions and repressive subcompartment like B2, and the transition domains are correlated with some intermediate subcompartments such as A2, B1 and B3, and usually locate in bivalent chromatin regions (**Fig. S16B**). Moreover, in combination with three types of boundary regions we defined, the components of the boundary types for different replication domains are shown in **Fig. 6C**. As the replication timings changed from early to late, the proportion of the WBs and NSBs for these domains gradually decrease, while the proportion of the NWBs significantly increase. This is consistent with our conclusion that DNA sequences near the WBs and NSBs replicated earlier, while sequences around the NWBs replicated late (**Fig. S12B**). Lastly, the major activities of the DNA replication within five types of replication domains occurred in different phases of the cell cycle, for example, the early replication domains mainly focused on G1 and S1 phases, the three types of transition domains corresponded to S2, S3, and S4 phases, while the replication of DNA sequences in the late replication domains concentrated in the G2 phase (**Fig. S16C**). Taken together, we give an example to show the strong relationship between replication domains, boundary types, epigenomic modifications, DNA replication timing and gene expression (**Fig. 6D**).

Moreover, the switch of replication types between domains in different cell lines might be accompanied with a shift in expression patterns of some essential genes. For example, we showed the distribution of ConsTADs and the corresponding replication types for the same region in GM12878 and K562 (**Fig. S17C**). In addition to the differences in topological domain configuration, the replication types of these domains also changed a lot, and the most significant domain-type reversals occurred near the genes *ITGA4* and *NCKAP1. ITGA4*, a homing receptor for lymphocytes, locates in an early replication domain in GM12878 and is accompanied by active modifications, thus exhibiting stronger gene expression. However, in K562, *ITGA4* resides in a late replication domain and its expression is also downregulated. This may be related to the specific activity of lymphocytes. In contrast, another gene *NCKAP1*, also known as *HEM-2*, is located near the boundary between two transition domains with relatively late replication timing, and shows low expression in GM12878. But *NCKAP1* resides near the boundary of two earlier replication domains in K562 and possesses some active makers like H3K4me3 and H3K36me3, thus showing significantly stronger gene expression. *NCKAP1* is known to correlate with several cancer types (59, 60), and another gene also belonging to the HEM family, HEM-1, is reported to be associated with the transition of fetal liver hematopoiesis to bone marrow (61). Therefore, the function of *NCKAP1* in K562 or leukemia may deserve further investigation. These results suggest that the topological domains as well as their replication types might serve as a bridge to understand some cell type-specific gene expression patterns.

Based on these normalized Repli-seq signals, the ConsTADs were also divided into five clusters with different relative replication patterns (**Fig. 6E**). We defined the direction of replication for each domain cluster according to the average signal profiles (**Fig. 6F**). For most domains, replication starts from the boundary at one side and then extends to the other side. Some domains replicate inward from both boundaries, but one side replicates significantly earlier, and only a small part of domains replicate from the inside to boundaries on both sides. These results are consistent with a previous study in which Petryk et al. found that replication origins in the human genome typically overlapped with TAD boundaries, while replication could terminate dispersedly between initiation zones (62). Moreover, the boundaries with earlier replication have stronger CTCF binding strength (**Fig. 6G**). These results suggest that the topological domains are the basic units of DNA replication and the direction of replication within domain is closely related to the CTCF binding strength around the boundary regions.

At last, we propose a novel model of replication domain (**Fig. 6H**). The topological domains serve as the basic units of DNA replication. We can divide them into early and late replication domains, and transition domains, which have stable and variable replication patterns, respectively. The direction of DNA replication within each domain is related to the CTCF profiles, and the boundary with stronger CTCF binding strength tends to replicate earlier. Furthermore, each type of replication domain has dominant epigenomic modifications, such as H3K27ac for early replication domains, H3K27me3 for transition domains, and H3K9me3 for late replication domains, implying that these domains not only have different DNA replication patterns but also belong to different chromatin states, including active, facultative and repressive, respectively.

## Discussion

In this study, we proposed a computational framework to integrate the TADs from 16 methods and constructed the TAD separation landscape based on their consensus. We demonstrated that the TAD separation landscape could not only accurately depict the locations of boundaries, but also served as an indicator to facilitate the comparison of topological structures across multiple cell lines. Moreover, we revealed three types of domain boundaries with different transcriptional activity, DNA replication timing and enrichment patterns of repeat elements like Alu and TcMar-Tigger. At last, we define the ConsTADs to represent the topological domains captured by the consensus of the 16 methods, which enhance our understanding of epigenomic patterns and chromatin state organization within topological domains.

Here, the three types of boundary regions could provide an explanation for the inconsistency among methods from the data perspective. Most methods could identify the NSBs and the bins they selected as the TAD boundary were highly fixed, while some methods could miss the NWBs due to their weaker insulation effect. Near the WBs, different methods might disagree on which bin to choose. However, the inconsistency among methods may also stem from their algorithmic principles and designs.

There are several directions for future studies. First, the TAD separation landscape could contribute to exploring the dynamics of chromatin organization across a series of time points or developmental stages in some biological processes such as the cell cycle or embryonic development. Second, the proposed strategy for defining TAD separation landscape and ConsTAD can be easily extended to more TAD-calling methods, facilitating the evaluation of some newly developed methods or integrating them to obtain more robust and consensual topological domains. Last, the three types boundaries could be used to explore the topological organization in the bulk Hi-C contact maps and those for individual cells. For example, we observe that the segments defined based on single cell 3D genome (2) corresponding to the three types of boundary regions also exhibit higher boundary probabilities than those labeled as non-boundaries (**Fig. S14A**). The segments labeled as WBs and NSBs could serve as domain boundaries in more cells and correspond to some apparent boundary structures in the proximity frequency matrix built from all single cells (**Fig. S14B**). These results reflected the high heterogeneity of topological domains at the single-cell level. The boundary regions we defined should correspond to the relatively conserved domain boundaries among single cells.

## Materials and Methods

### Hi-C data

In this study, we used *in situ* Hi-C data for seven human cell lines, including GM12878, HMEC, HUVEC, IMR90, K562, KBM7 and NHEK (GEO accession: GSE63525). GM12878 contains two replicates with DpnII and MboI as the restriction enzyme, respectively, and the rest of cell lines only have one with MboI as the restriction enzyme. Comparison and integration of 16 TAD-calling methods were performed based on the intra-chromosomal contact maps generated at 50kb resolution for chromosome 2 of these seven cell lines.

### 3D chromatin imaging data

We downloaded the file containing 3D coordinates of the genomic loci across chromosome 2 of IMR90 single cells measured using the sequential hybridization approach (2) from Zenodo (https://zenodo.org/record/3928890).

### ChIP-seq, DNase-seq, Repli-seq, RNA-seq, chromatin state, subcompartment, repeat elements and other biological data

To compare and assess the 16 TAD-calling methods and explain the biological significance of TAD separation landscape, we collected the following biological datasets.

First, we obtained some ChIP-seq datasets. For CTCF, the ChIP-seq narrow-peak datasets for GM12878, HMEC, HUVEC, IMR90, K562, NHEK were available at ENCODE Uniform TFBS composite track from the UCSC genome browser (65): wgEncodeEH000029, wgEncodeEH000419, wgEncodeEH000551, wgEncodeEH002831, wgEncodeEH000042, wgEncodeEH000406. For RAD21 and SMC3, we downloaded the ChIP-seq narrow-peak files for GM12878 (accessions: wgEncodeEH001640 and wgEncodeEH001833). Besides, we also collected several ChIP-seq datasets in bigwig format for some TFs like CTCF, RAD21, SMC3, Pol2 and YY1, as well as some histone modification marks including H2A.Z, H3K27me3, H3K9me3, H3K27ac, H3K4me1, H3K4me2, H3K4me3, H3K9ac, H3K36me3, H3K79me2, H4K20me1 for GM12878 and K562. As for DNase-seq, we obtained the narrow-peak files for DNase accessibility sites (wgEncodeAwgDnaseUwdukeGm12878UniPk.narrowPeak.gz and wgEncodeAwgDnaseUwdukeK562UniPk.narrowPeak.gz) and track signals in bigwig format for GM12878 and K562 from the UCSC genome browser. All these files in bigwig format can be download from the website: (http://ftp.ebi.ac.uk/pub/databases/ensembl/encode/integration_data_jan2011/byData_Type/signal/jan2011/bigwig).

For DNA methylation, we obtained the MeDIP-seq data in bigwig format for GM12878 and K562 (GEO accessions: GSM1368907 and GSM1368906).

For Repli-seq data, we downloaded the processed track signals belonging to the six phases of cell cycle: G1, S1, S2, S3, S4, and G2 as well as the wavelet-smoothed signals for GM12878 and K562 (GEO accessions: GSM923451 and GSM923448).

We also obtained uniformly processed RNA-seq data including the stranded track signals in bigwig format as well as the RPKM expression matrices for protein coding genes in GM12878 and K562 from the Roadmap consortium (66).

As for chromatin state, we collected the ChromHMM 15-state results (45) for GM12878, HMEC, HUVEC, IMR90, K562, NHEK from the Roadmap consortium (https://egg2.wustl.edu/roadmap/web_portal/chr_state_learning.html). We also obtained the Segway 5-state results (51) for GM12878, HUVEC, IMR90, K562 from the ENCODE consortium (67, 68): ENCFF058KWZ, ENCFF202BIE, ENCFF762DWI, ENCFF996BDE.

As for the subcompartment annotation, we downloaded the subcomaprtments predicted by SNIPER (52) for GM12878, HMEC, HUVEC, IMR90, K562, including five kinds of subcompartments: A1, A2, B1, B2, B3.

We downloaded the repeat elements from the UCSC Genome Browser (http://hgdownload.cse.ucsc.edu/goldenPath/hg19/database/rmsk.txt.gz), and the phastCons scores for chromosome 2 of human genome assembly hg19 (https://hgdownload.cse.ucsc.edu/goldenpath/hg19/phastCons46way/vertebrate/) to indicate the sequence conservation between human and other species. We obtained the list of human housekeeping genes from (69).

### TAD-calling methods

Since the concept of TAD was introduced, a number of TAD-calling methods have been developed. We collected 16 TAD-calling methods for comparison and integration, including some classical ones, such as DI (9) and IS (10), as well as methods like TopDom (25), which was considered to perform well in the tests of Zufferey et al. (23). But the methods like TADtree (70), TADbit (71) and matryoshka (72) running too slow, and some methods like Arrowhead (7) with rare requirements on the input data format were excluded from this study. Besides, we also introduced some newly developed methods, such as deDoc (37) and MSTD (29).

We tried to utilize these methods with default parameters or set these parameters according to the authors’ recommendations for their packages (**Table S1**). In addition, when the method requires the input Hi-C data to be normalized, we uniformly used the iterative correction and eigenvector decomposition method (ICE) (73) for normalization. Then the outputs of different TAD-calling methods were converted to two files with uniform format, for recording the chromatin ranges of the topological domains and the locations of the start and end boundaries, respectively. Of note, for some methods like HiTAD and 3DNetMod outputting nested or overlapping domains, we only extracted domains with the lowest level, such that no further nesting contained in them.

### Comparison of different TAD-calling methods

We compared TADs identified by 16 TAD-calling methods on seven human cell lines by calculating some metrics between domain sets or boundary sets, such as the absolute differences of TAD number and average size, Jaccard index and the Measure of Concordance.

#### Jaccard index between boundary sets

Here we introduced a modified Jaccard index to evaluate the similarity between two sets of TAD boundaries A and B:

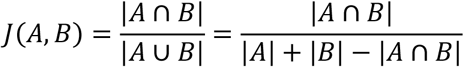

where 0 ≤ *J*(*A,B*) ≤ 1. Here, we adopt a tolerance radius of 1 bin when defining the intersection between two sets of TAD boundaries. This means that, if the distance between two boundaries doesn’t exceed 50kb, they will be determined as the shared ones between boundary sets.

#### Measure of Concordance (MoC) between domain sets

MoC was introduced by Zufferey et al. (23) to compare TAD partitions and it is defined as follows:

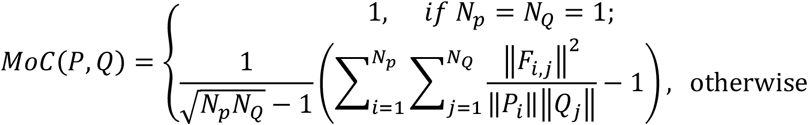

where *P* and *Q* are two domain sets, *N_p_* and *N_Q_* represent the number of TADs contained in these two sets. *P_i_* and *Q_j_* are two TADs with domain lengths of ||*P_i_*|| and ||*Q_j_*||, where ||*F_i,j_*|| denotes the overlap length between *P_i_* and *Q_i_*. Here the length indicates the number of base pairs on the DNA sequence.

We calculated these metrics mainly for comparison of TADs across methods or across datasets. For comparison across methods, we kept the dataset the same and calculated the metrics between TADs identified by different methods. For comparison across datasets, we kept the method the same and calculated the metrics between TADs found by this method on different Hi-C datasets.

#### Enrichment of structural proteins at TAD boundaries

For the structural proteins (e.g., CTCF, RAD21, and SMC) that have been reported related with TAD boundary, we counted the their binding peaks in each 10kb bin and built up the profile of median peaks around the boundaries identified by certain method. Then we computed the fold change between their binding peaks at TAD boundaries versus adjacent flanking regions.

### Boundary voting strategy

We integrated the results of different TAD-calling methods by a strategy of boundary voting. In this process, we firstly extract the boundary sets of each method and divide these boundaries into start boundaries and end boundaries. Then each bin identified as a boundary can contribute one score to the bins nearby according to a preset radius. In this study, we use the strictest radius of zero, which means the boundary could only contribute one score to itself. Thus, we can get two boundary score profiles from start and end boundaries, respectively. Then we take the profile of maximum value between these two boundary score profiles to ensure that a method contributes at most one boundary score to a bin. Finally, the profiles of boundary scores obtained by all these methods are summed, thus a boundary score is assigned to each bin on the genome, representing the number of methods that define it to be the boundary of a TAD.

According to the boundary score, we divide the TAD boundaries identified by each method into different intervals and show the proportions of the boundaries in each interval (**Fig. 2A**). If a bin with non-zero score does not belong to the boundary sets of certain method, we call this bin the missed boundary for that method, and conversely, the bin is called captured boundary. Thus, we exhibit the scores of boundaries captured or missed by certain methods (**Fig. 2B**). Moreover, for each method, we explore the relationship between the boundary score of missed boundaries and the minimum distance between that missed boundary and other captured boundaries (**Fig. S2C**).

### Assessment of the reliability for boundaries with different scores

The bins with non-zero boundary scores formed the collection of candidate boundaries of all 16 TAD-calling methods. These boundaries were divided into five intervals with different boundary scores. We then adopted three kinds of 1D topological indicators including the directionality index (DI), insulation score (IS) and contrast index (CI), as well as three kinds of structural biological markers including CTCF, RAD21 and SMC3 to assess the reliability of these boundaries in different score intervals. These three 1D topological indicators are computed according to Fig. S2D. They have been used by some methods to identify TAD boundaries from Hi-C interaction maps, and they show distinct signal patterns at the TAD boundaries, reflecting the isolation strength of the boundaries. While these structural proteins tend to enrich at TAD boundaries and can exhibit peaks at boundaries in their binding profiles. Here, we showed the profiles of three 1D topological indicators and three structural proteins for boundaries in each score interval.

### Distribution plot of the scores for two boundaries of TADs

For each TAD-calling method, we collected the scores for two boundaries of each TAD and estimated the density profiles for these paired boundary scores with a Gaussian kernel. We applied the function *seaborn.jointplot* in a python package named seaborn to achieve it and to draw the density plots.

### Construction of TAD separation landscape

Based on the boundary voting strategy, we got the original boundary score profile for bins along the genome. However, these original profiles contained many bins with very low scores, probably leading to some unreliable boundary locations (**Fig. S6E**).

Moreover, if we considered consecutive bins with non-zero scores as possible boundary regions, these unreliable boundary locations could make the adjacent boundaries too close to each other, which was not consistent with the perception of TAD length in previous study (**Fig. S6F**). Hence, we introduced the contrast *p*-value to refine the original boundary score profile.

Firstly, we applied a distance-dependent z-score normalization to the Hi-C contact map to alleviate the effect of genomic distance on the strength of the chromatin interactions. In this process, according to the resolution of contact map, we calculated the z-score for the chromatin interactions at each distance (**Fig. S5A**). Then we extracted the normalized interactions of two kinds of regions, including the upstream and downstream regions as well as the insulation regions, to calculate the contrast *p*-value for each bin along the genome (**Fig. 3A**). We used a single-side Mann-Whitney U test to test whether the insulation region showed weaker interactions compared with upstream and downstream regions, which would output the contrast *p*-value. A very small contrast *p*-value reflects a strong insulation of chromatin interactions at the current bin. We presented different window sizes for the calculation of contrast *p*-value and selected the most suitable one according to the Pearson correlation between the original boundary score profile and the contrast *p*-value profiles with different windows (**Fig. S5B**). We then used the profile of contrast *p*-value as a reference and refined the original boundary score profile by three kinds of operations, including **Add**, **Filter** and **Combine**. The **Add** operation will add 1 score to bins with zero boundary scores but with contrast *p*-values below a preset cut-off. The **Filter** operation will turn the boundary scores to zero for bins with *p*-values greater than the cut-off, but for bins in the valleys of *p*-value profiles, the boundary scores are kept. The **Combine** operation will combine two adjacent boundary regions separated by one bin gap and the gap will be filled with the average boundary score of the upper and lower bins (**Fig. S5C**). In this study, we set a cut-off of 0.05 for contrast *p*-value. The refined boundary score profile reflected the boundary regions with higher score and the distances between adjacent boundary regions were more reasonable (**Fig. S5E-J**). We termed the refined boundary score profile as the TAD separation landscape, which could accurately depict the regions of TAD boundary and were highly consistent with the domain patterns in Hi-C contact maps (**Fig. S5D**).

### Analyses of TAD separation landscapes of multiple cell lines

We built the TAD separation landscapes of seven human cell lines, including GM12878, HMEC, HUVEC, IMR90, K562, KBM7 and NHEK, combined them with other data to explore their biological significance, and used them as references for deciphering conserved and cell type-specific boundary regions among multiple cell lines.

For each bin along the genome, we calculated the average boundary scores among seven cell lines based on their TAD separation landscapes. We sorted these bins by the average scores in ascending order, and selected some quartiles, including 0%, 30%, 44%, 58%, 72%, 86% and 100%, to divide all bins into six levels. The bins contained in the first level (0~30%) all have a boundary score of 0 and bins with higher levels would have larger boundary scores. For bins with different levels, we computed the number of cell lines in which they have non-zero boundary scores and counted the number of housekeeping genes in each bin, as well as the average number of CTCF binding peaks across cell lines, and we also calculated the average phastCons scores for DNA sequences of these CTCF binding peaks. Besides, we clustered the Hi-C samples from different cell lines based on the Pearson correlations between their TAD separation landscapes.

We then performed the comparison of TAD separation landscapes calculated for different cell lines. For pairwise comparison of cell lines, we firstly identified the core boundary regions between them. We added up the TAD separation landscapes of the two cell lines and used a cut-off to filter out bins with scores below two, which meant that the scores of these bins were converted to zero in the added score profile. We defined the consecutive bins with non-zero scores in the added score profile as core boundary regions between them For each core boundary region, we examined the maximum score of the corresponding region in the TAD separation landscapes of the two cell lines. If the maximum scores of core regions in the two cell lines are both above five, we defined these regions as conserved boundary regions. For a core boundary region, if the maximum score exceeds five only in one cell line, while the maximum score does not exceed two in the other one, and the difference between these two maximum scores is equal to or greater than five, then we consider it as a cell type-specific boundary region. This boundary is gained in one cell line and lost in the other. For multiple comparison of cell lines, we also added up the TAD separation landscapes and filtered bins with a cut-off of five to identify the core boundary regions. If the maximum score of the corresponding region was larger than five in the TAD separation landscape of a certain cell line, we consider this cell line possesses this core boundary region. Thus, we can identify the boundary regions which are conserved in all seven cell lines, specifically occurred in a certain cell line (**Fig. S7E and Fig. S8**), or shared by several cell lines (**Fig. S9**).

### Drawing aggregated Hi-C contact maps

We separately normalized the Hi-C contact matrices of different cell lines by dividing the maximum value in them (**Figs. 4A, 4H, 4I and Fig. S9**). To compare the Hi-C contact maps around a set of boundary regions, we firstly normalized them by dividing the maximum value and then computed the average contact maps for demonstration. Besides, the average boundary score profile around these boundary regions were also shown below the contact maps (**Figs. 4D, 4F, 4G and Fig. S8**).

### Identification of three types of boundary regions

We defined the consecutive bins with non-zero scores in the TAD separation landscape as boundary regions for seven cell lines. For each cell line, the length and average score of each boundary region are used as features and then the boundary regions are clustered using K-means to select the optimal cluster number according to the within-cluster sum of squared error and silhouette coefficient.

The within-cluster sum of squared error (WCSSE) is the sum of the squared differences between each sample and its cluster center across all cluster:

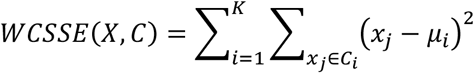

where *K* denotes the cluster number, *X* = {*x*_1_, *x*_2_, …, *x_n_*} denotes the set of boundary regions, *C_i_* is the *i*-th cluster of K-means and *μ_i_* is the cluster center. The number of clusters that minimize the sum of squared error can be viewed as optimal.

The silhouette coefficient (SC) is a measure of how similar a sample is to its own cluster compared to other clusters, i.e.,

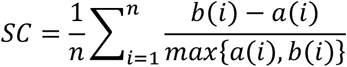

where 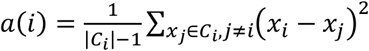 reflects the average distance for sample *i* within a cluster and 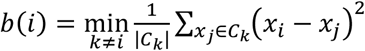 reflects the minimum average distance between sample in other cluster for sample *i*. The number of clusters that maximize the silhouette coefficient can be viewed as the optimal one.

According to the within-cluster sum of squared error and silhouette coefficient, we found that the optimal number of clusters of K-means for all the cell lines was three. Then under the optimal number of clusters, two thresholds for the length and the average score of boundary region are selected based on the clustering results of K-means. Based on these thresholds, all boundary regions are classified into three types: the narrow-strong boundary regions (NSBs), narrow-weak boundary regions (NWBs) and wide boundary regions (WBs) (**Fig. S10**).

### Analyses of three types of boundary regions identified in multiple cell lines

We identified three types of boundary regions in seven cell lines and selected GM12878 and K562 as representatives to explore the biological significance of these boundary regions. We profiled three 1D topological indicators and three structural proteins’ binding peaks around different types of boundary regions. For topological indicators, the values in each 50kb bins are shown. For structural proteins, we counted their binding peaks in each 50kb bin and computed the average peaks in all bins along the chromosome as the background, we then calculated the fold change by dividing the background from peak number in each bin (**Figs. 5E and 5F)**.

We then explored the cross cell-type conservation of three types of boundary regions. If one boundary region in a cell line overlaps with the one in another cell line, and the region type remains the same, we consider it to be conserved between these two cell lines. Hence, for each boundary region, we computed and displayed the ratio of cell lines in which it was considered as conserved (**Fig. 5D**).

We collected the track signals in bigwig format for multiple biological data including histone modifications, TF binding, DNA methylation, chromatin accessibility, gene expression and DNA replication timing to explore their enrichment in three types of boundary regions. For each kind of data, we firstly computed the average signals of it in each 50kb bin along the chromosome and selected the median values of these average signals as the background. We then calculated the average signal in each boundary region and extracted the median of the average signals of all ones belonging to the same type. At last, the fold change was obtained by dividing the median values across the chromosome (**Fig. 5G and Fig. S14D**).

As for the Repli-seq data for six phases of cell cycle, we got the average values in all 50kb bins along the chromosome and used the mean value of them as the background. Then the values in each bin were divided by the background to get the fold change. Finally, the profile of these fold change around three types of boundary regions were displayed according to the phase order in cell cycle (**Fig. 5H and Fig. S14E**). The wavelet-smoothed signals are weighted average of tracks of these six phases, in which higher values correspond to earlier replication. Thus, we also showed the profile of fold change of wavelet-smoothed signals around different types of boundary regions (**Fig. S13B and Fig. S14F**).

We also performed the enrichment analysis of chromatin states and subcompartments in different types of boundary regions. For certain type of chromatin state or subcompartment, the fold change (FC) of enrichment was defined by dividing the length of a boundary region belonging to it by the expected length. The expected length was calculated according to the total length of boundary regions and the distribution of chromatin state or subcompartment on the chromosome:

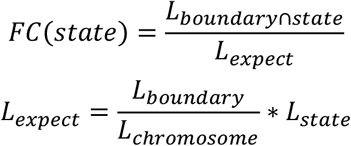

where the *L_boundary⋂state_* denotes the length of overlap between a boundary region and a chromatin state or subcompantment, *L_boundary_*, *L_state_* and *L_chromosome_* represent the total lengths of boundary regions, chromatin state and chromosome, respectively.

We explored the enrichment of some repeat elements in three types of boundary regions. These repeat elements mainly consist of seven classes, such as the short interspersed nuclear elements (SINE), the long interspersed nuclear elements (LINE) and the DNA repeat elements, and each class also contains some subclasses. For each boundary region, we randomly selected 500 regions on the chromosome of equal length to the boundary region, and counted the number of repeat elements located in the boundary region and these random regions, respectively. We then calculated the enrichment z-score of repeat elements in the boundary region based on the mean and variance of the elements in random regions. Hence, a z-score above one indicates the repeat element is enriched in boundary region (**Fig. 5J and Fig. S12**).

We then compared the TFs enriched in different types of boundary regions. First, we collected the accessible sections located in boundary regions based on DNase-seq data and extracted their DNA sequences. Then we used HOMER (56) to find the motifs of TFs in these DNA sequences and calculated the significance of their enrichment with the hypergeometric test. TFs were then ranked by descending negative log(*p*-value) and the top 25 TFs for each type of boundary region were selected for comparison (**Figs. S13C and S13D**).

### Exploring boundary probabilities of boundary regions at single cell level

We obtained the 3D coordinates for probes representing 250kb loci on chromosome 2 of IMR90 single cells and these coordinates are separated for each imaged chromosomal copy. We first calculated the 3D spatial distances between pairs of imaged chromatin loci and constructed the spatial distance matrices for every single chromosome. We then identified chromatin domains in single chromosomes following the procedures proposed by Su et al. (2022). In brief, a modified insulation score based on the spatial distance matrices was calculated and the domain boundaries were recognized as the peaks in the profile of insulation score (2). In this way, we can assign a boundary probability to each probe, indicating the number of single chromosomes that consider it as the domain boundary over the total number of chromosomes. We then identified the boundary regions on chromosome 2 of IMR90 with 50kb-resolution Hi-C contact map and used the LiftOver (74) to convert all 50kb bins from human genome assembly hg19 to bins on hg38 assembly. For each probe representing a 250kb locus, we assigned it a label corresponding to the type of boundary region that covered the largest proportion of it, and if a probe did not overlap with any boundary regions, it would be defined as non-boundary. Besides, for each single chromosome, if the distance between two loci is below 500nm, they will be considered as contacting with each other. Thus, we got the overall proximity frequency matrix of all these single chromosomes by dividing the contact frequency of each pair of loci by the total number of single chromosomes (**Fig. S14**).

### Define the ConsTAD by a boundary matching strategy

Based on the TAD separation landscape, we proposed a boundary matching strategy to select two bins from the adjacent boundary regions and the intermediate regions between these bins form a ConsTAD. For each pair of adjacent boundary regions, we try all paired combinations of these bins and select the best one by calculating the average boundary score of the two selected bins and the fold change of the contacts within the domain relative to the upstream and downstream background regions (**Fig. S15A**). In detail, we firstly applied a distance-dependent z-score normalization to the Hi-C contact map and extracted the domain region according to the two selected bins as well as the upstream and downstream regions according to the distances between current boundary regions and the flanking ones on each side. We then calculated the fold change of average signal within a domain relative to the average signal within upstream and downstream regions. Thus, we could get the rank of fold change as well as the rank of average boundary score among all paired combinations of bins, the combination with the best average rank in terms of the two aspects would be selected to form the ConsTADs.

### Analysis of the patterns of epigenomics signals within ConsTADs

We identified the ConsTADs on chromosome 2 in GM12878. For each domain, we extended 20% of the domain length upstream and downstream respectively. Then the domains together with their flanking areas were divided into 700 intervals with equal length and the mean epigenetic signal in each interval was calculated. For each domain, the 700-dimensional epigenetic signal vector was scaled into [0, 1] by the min-max normalization. We then calculated the average epigenetic signal profile across all ConsTADs and applied the function *scipy.signal.savgol_filter* in a python package called scipy to smooth the profiles for visualization (**Fig. 6A**).

### Identification of topological domains with significant H3K36me3/H3K27me3 differential signal

We calculated the average signal of H3K36me3 and H3K27me3 in each 50kb bins along the chromosome 2 in GM12878 and then got the fold change of signal for each bin by dividing the mean signal across the whole chromosome. For each bin, we computed the log2-ratio between the H3K36me3 and H3K27me3 fold change, similarly to Zufferey et al. (23), we termed them as LR values or LR intervals and a positive value indicates a bias to H3K36me3, while a negative one indicates a bias to H3K27me3, and these biases usually stay the same in some consecutive intervals and the positions with altered bias are recorded. We collected the topological domains identified by all 16 TAD-calling methods as well as the ConsTADs we defined and calculated the distances from all bias-changing points to the nearest domain boundary for each method (**Fig. S15C**). We also calculated the average LR values within each domain and shuffled the LR intervals 1000 times to derive a null distribution of LR values within domains. For each domain, an empirical *p*-value can be calculated by comparing its observed average LR values with the null distribution. The *p*-values for all the domains identified by each TAD-calling method were corrected using the Benjamini-Hochberg procedure and domains with a corrected *p*-value smaller than 0.1 were considered with significant H3K36me3/H3K27me3 differential signal. Domains under 150kb (three bins at 50kb resolution) in length were excluded from this analysis because they were too short (**Fig. 6B**).

### Clustering topological domains based on DNA replication signal

We collected topological domains identified by 16 TAD-calling methods and the ConsTADs for chromosome 2 of GM12878. Each domain was divided into 500 equal-length intervals and the average wavelet-smoothed signals of Repli-seq for each interval was also calculated. Then an agglomerative clustering was applied to each domain sets based on Euclidean distance and ward’s method (**Fig. S15D**). For each method, the topological domains were divided into five clusters differing in the periods of DNA replication and bins within these domains were also assigned to certain cluster. Then for each bin along the chromosome, a consistency score of domain replication cluster was calculated as the proportion of methods in which the bin having the same cluster assignment as the current method (**Fig. S15F and G**).

We also obtained the relative DNA replication signal within each domain by using the min-max normalization to show the relative early or late of DNA replication among regions inside the domain. We then applied the agglomerative clustering to these relative DNA replication signals of ConsTADs and got five clusters with distinct relative replication modes (**Fig. 6E**).

### Enrichment analysis of chromatin states and subcompartments in replication domains

For each type of replication domains, we calculated the fold change of enrichment for chromatin states or subcompartments by dividing the observed state or subcompartment lengths in domains by the average ones across the chromosome.

## Code availability

https://github.com/zhanglabtools/ConsTADs

## Acknowledgements

This work has been supported by the National Key Research and Development Program of China [2019YFA0709501], the National Natural Science Foundation of China [62173271, 61873202, 61621003], the Strategic Priority Research Program of the Chinese Academy of Sciences (CAS) [XDPB17], and the Key-Area Research and Development of Guangdong Province [2020B1111190001].

